# A Mixture Model Incorporating Individual Heterogeneity in Human Lifetimes

**DOI:** 10.1101/2021.01.29.428902

**Authors:** Fei Huang, Ross Maller, Brandon Milholland, Xu Ning

## Abstract

Analysis of some extensive individual-record data using a demographically informed model suggests constructing a general population model in which the lifetime of a person, beyond a certain threshold age, follows an extreme value distribution with a finite upper bound, and with that upper bound randomized over the population. The resulting population model incorporates heterogeneity in life-lengths, with lifetimes being finite individually, but with extremely long lifespans having negligible probability. Our findings are compared in detail with those of related studies in the literature, and used to reconcile contradictions between previous studies of extreme longevity. While being consistent with currently reported analyses of human lifetimes, we nevertheless differ with those who conclude in favour of unbounded human lifetimes.

---

The question of whether or not a limit to human lifespan exists may be as old as civilization itself. Earliest speculation was hampered by a lack of reliable age data, biological knowledge and statistical methods. Foundations for the modern field of demography were laid in the 19^*th*^ Century by Gompertz, who modelled mortality as increasing exponentially with age^1^ and Makeham, who added an age-independent term to the equation^2^. The Gompertz-Makeham Law of Mortality now enjoys near-universal acceptance as an accurate model for mortality at most ages. However, scientists still fiercely debate whether mortality deviates from the Gompertz-Makeham law at very old ages and whether human longevity can be increased into the future. On these issues, there is no consensus, with some forseeing many years of extended longevity^3^, while others argue that previous upward trends are largely played out and only modest improvements can be expected in the future^4^.

Dong, Milholland & Vijg^5^ analysed trends in three public longevity datasets: demographic data in 41 countries and territories from 1900 onwards contained in the Human Mortality Database (HMD)^6^; the yearly maximum reported ages at death of individuals aged 110 years or more (“supercentenarians”) from France, Japan, the UK and the US available in the International Database on Longevity (IDL)^7^, a total of 534 individuals over the years 1968–2006; and maximum reported ages at death reported by the Gerontology Research Group (GRG)^8^ for the years 1955–2015. They deduced a continuing increase in human life expectancy since 1900, but with a rate of improvement in survival which peaks and then declines for very old ages. Linear regression techniques were used to show that the increase in ages at death halted at around 1995. The time series of the estimates of the age with greatest improvement in survival appeared to level off around 1980, rather than continuing to increase with calendar year. It was inferred from this that human lifespan may have a natural limit. Some lively commentary which followed on publication of that paper is summarised later in the present paper.

Huang, Maller and Ning^9^ analysed a high-quality Netherlands dataset, based on the Human Mortality Database^10^ (HMD), augmented with a dataset from the Centraal Bureau voor de Statistiek of the Netherlands, to construct lifetables separately for females and males aged 65+ born in 1-year cohorts from 1893 to 1908. The data extends to 112 years of age for females and 110 years for males (maxima over all cohorts) and contains a substantial number of individuals aged 95 years or more at death. Data on a total of 304,917 individuals was available. The lifetables were constructed by fitting to each cohort a model consisting of a traditional Gompertz distribution up till a data determined threshold age *N*, followed by an extreme value distribution for ages greater than *N*. A constraint was imposed so that the hazards in the two parts of the model transitioned smoothly between them.

The mathematical formulation of the extreme value part of the model contains a parameter, γ, which can be used to judge whether the right extreme of the fitted distribution is finite or not. This effectively constitutes a test for whether the data indicates a finite or unbounded lifetime for the individuals in the cohort. A negative estimated γ indicates a finite upper bound, whereas a positive γ denotes an unbounded distribution.

In each of the 16 cohorts of the Netherlands data analysed^9^, and for both females and males, the estimated γ was significantly negative (Table 1). So the data and the model strongly suggest a finite upper limit to individuals’ lifetimes. This limit, however, varies over a range of around 110-118 years for the different cohorts, with standard errors of the order of 2-4 years (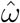 in Table 1).

**Table 1:** Estimated parameters for each birth cohort (reproduced from^9^). Females: 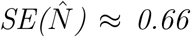 years, 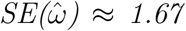 years. Males: 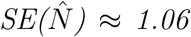 years, 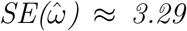 years. (Standard errors averaged over cohorts.) *6 outlying extreme values omitted for males 1907 analysis.

|  | Females |  |  | Males* |  |  |
| --- | --- | --- | --- | --- | --- | --- |
| Cohort | $\hat{\gamma}$ | $\hat{\omega}$ | $\hat{N}$ | $\hat{\gamma}$ | $\hat{\omega}$ | $\hat{N}$ |
| 1893 | -0.159 | 114.74 | 97 | -0.221 | 109.55 | 89 |
| 1894 | -0.160 | 114.13 | 98 | -0.195 | 110.42 | 91 |
| 1895 | -0.205 | 110.67 | 98 | -0.221 | 109.00 | 90 |
| 1896 | -0.173 | 112.95 | 98 | -0.176 | 110.38 | 98 |
| 1897 | -0.207 | 110.88 | 97 | -0.179 | 110.37 | 98 |
| 1898 | -0.154 | 114.77 | 98 | -0.151 | 112.27 | 98 |
| 1899 | -0.171 | 113.10 | 98 | -0.179 | 110.82 | 95 |
| 1900 | -0.208 | 110.24 | 98 | -0.134 | 115.17 | 96 |
| 1901 | -0.191 | 111.78 | 97 | -0.181 | 111.12 | 94 |
| 1902 | -0.151 | 114.78 | 98 | -0.109 | 118.56 | 97 |
| 1903 | -0.133 | 116.59 | 98 | -0.116 | 117.01 | 98 |
| 1904 | -0.174 | 113.52 | 96 | -0.197 | 109.65 | 93 |
| 1905 | -0.144 | 116.03 | 97 | -0.139 | 113.87 | 98 |
| 1906 | -0.135 | 116.02 | 98 | -0.125 | 117.31 | 95 |
| 1907 | -0.166 | 113.35 | 97 | -0.145 | 112.92 | 98 |
| 1908 | -0.208 | 111.11 | 95 | -0.134 | 114.94 | 96 |

Based on these findings and those of Dong et al.^5^, we are led to formulate a model which is consistent with there being a finite lifetime for each person but which does not impose an absolute finite upper limit to human lifetimes overall. We suggest that with each person is associated a (negative) gamma parameter, signaling a finite lifetime for that person, which is specific to the person, but varies over the population according to some distribution. This is done by introducing a mixing measure to randomize the lifetime upper limit for each person.

In the next two sections we show how these ideas can be implemented. In later sections we review published discussions relating to the possible existence of an upper limit to human lifetimes, and discuss how advanced age mortality “acceleration and deceleration” effects are included in the Huang et al.^9^ model, and how the “three laws of biodemography” are satisfied by it.

## Advanced age life table modeling

We briefly recap the extreme value theory approach adopted to analyse the Netherlands data^9^. We reproduce in Figure 1 the empirical mortality rates (conditional probability of dying in the next year, given survival to the designated age) for the complete dataset (231,129 females and 73,788 males, totaling 304,917 persons).

**Figure 1:**
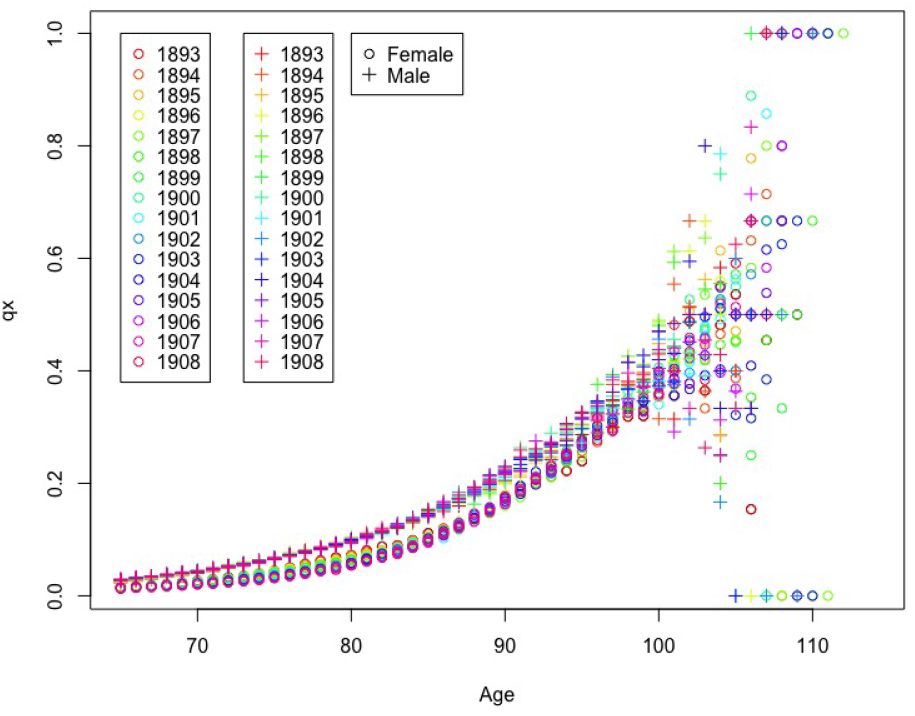
*Observed Netherlands Death Probabilities*

To this data was fitted a model consisting of a traditional Gompertz distribution up till a data determined threshold age *N*, followed by an extreme value distribution (a generalised Pareto distribution, GPD) for ages greater than *N*. A person’s age-at-death is then given by *T* = *N* + *Y*, where *Y* is a continuous random variable having cumulative distribution function (cdf)

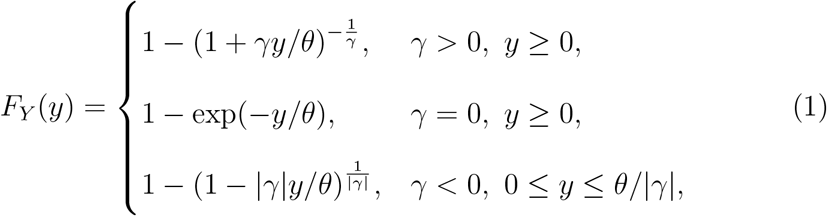

and *θ* > 0 and 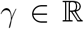 are constants. An innovative feature of the model, not present in earlier models, is that a smooth transition in hazard rates between ages less than and greater than the threshold age *N* is enforced. The resulting “smooth threshold life table” (STLT) model^9^ was fitted to the Netherlands data by maximum likelihood, producing the estimates of γ and *θ* in Table 1 for each sex × year-of-birth cohort.

We confine attention here to the extreme upper range of the data. When γ ≥ 0 in (1), the distribution has an infinite right endpoint and indefinitely large lifetime ages are potentially possible, with γ = 0 corresponding in particular to an exponential distribution. But when γ < 0, the distribution has a finite right end point at *θ*/|*γ*| corresponding to an upper bound on a lifetime at age N+*θ*/|*γ*|; ages greater than this cannot occur (have probability 0) in this model. Thus the estimated value, 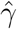, of *γ* from a dataset is relevant to the question of whether there is a finite upper limit to the lifetime of an individual from the population (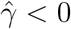) or no such upper limit (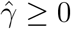).

The estimates for *γ* in Table 1 indicate for all cohorts a finite upper limit to the distribution of individuals’ lifetimes. This is suggestive of a finite lifetime for a person. But we do not conclude from this the existence of a “hard” upper limit to human lifetimes; the estimated upper limits vary with cohort and gender, and, potentially, from person to person. As a way of allowing for this, we randomize the upper limit by introducing a mixing measure for each person.

## A mixture model for lifetimes

We write the lifetime *T* of a person in a given population as *T* = *N* + *Y*, where *N* is the threshold age at which the extreme value distribution takes effect, and Y has the distribution in (1) corresponding to a parameter value of *γ* < 0:

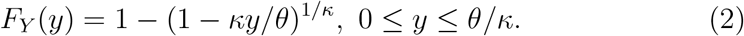

(We set *κ* = |*γ*| > 0 to avoid many modulus signs.) Now assume that associated with person *i* is a positive random variable *V_i_*, so that *Y_i_* for person *i* has the distribution in (2), conditional on the given value of *V_i_*:

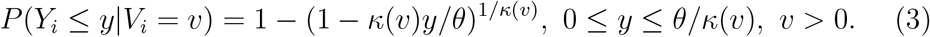

Here the constant *κ* is replaced by *κ*(*v*), a positive function of *v* to be specified.

Differentiating (3) gives the conditional density of *Y_i_* as

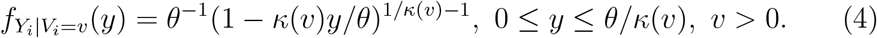

This model has features in common with the concept of “frailty” introduced in mortality modeling by Vaupel, Manton & Stallard^11^, which has subsequently found great application in lifetime survival analysis. In their context, the heterogeneity is assumed to apply to the probability or hazard of dying at a particular age, whereas in ours it refers to the life-length. The frailty concept allows for random unobserved factors. Those with high “frailty” tend to die earlier, leaving those with lower frailty. By comparison, in our model individuals with a lower life-length (analogous to a higher innate frailty) die sooner, while those with a higher individual life-length (analogous to lower frailty) survive longer. Common assumptions in frailty analysis are that the corresponding *V_i_* have a lognormal or gamma distribution.

To illustrate the possibilities, suppose in our setup that the *V_i_* have a Gamma(1/*α*,1/*α*) distribution for some *α* > 0. This parametrization of the gamma distribution implies a scaling such that *E*(*V_i_*) = 1. (The mixing variable is only identifiable up to a scale factor. We take the value 1 as specifying the distribution of a reference individual.) Consequently, overall, the (*Y_i_*) are taken to be independent and identically distributed with density

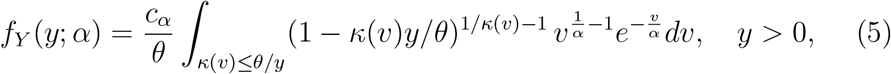

where *c_α_* = 1/α^1/α^Γ(1/α). The excess lifetime *Y* may take values in *y* ∈(0, ∞) hence is unrestricted but with probability 1 takes only finite values.

The choice of the function *κ*(*v*) is at our discretion and can be made optimally in a particular sample. A simple choice is to set *κ*(*v*) = *κ × v^β^*, *v* > 0, where *κ* and *β* are constants to be estimated. In Fig. 2 we take *N*, *κ* and *θ* as the estimated values for the 1893 female cohort in the Netherlands data, and illustrate the range of distributions possible for lifetimes *T* inherent in (5) by plotting *f_Y+N_*(*y + N*; *α*) for *β* = 7 and various values of *α* > 0. The densities are quite exponential-like, which is to be expected since the underlying density in (4) tends to an exponential density as *κ* → 0, hence should be close to exponential for small values of *κ*, as we have in Table 1. The densities in Figure 2 assign negligible probabilities to values of *N + Y*(= *T*) greater than 115 years or so, though this probability is never exactly 0.

**Figure 2:**
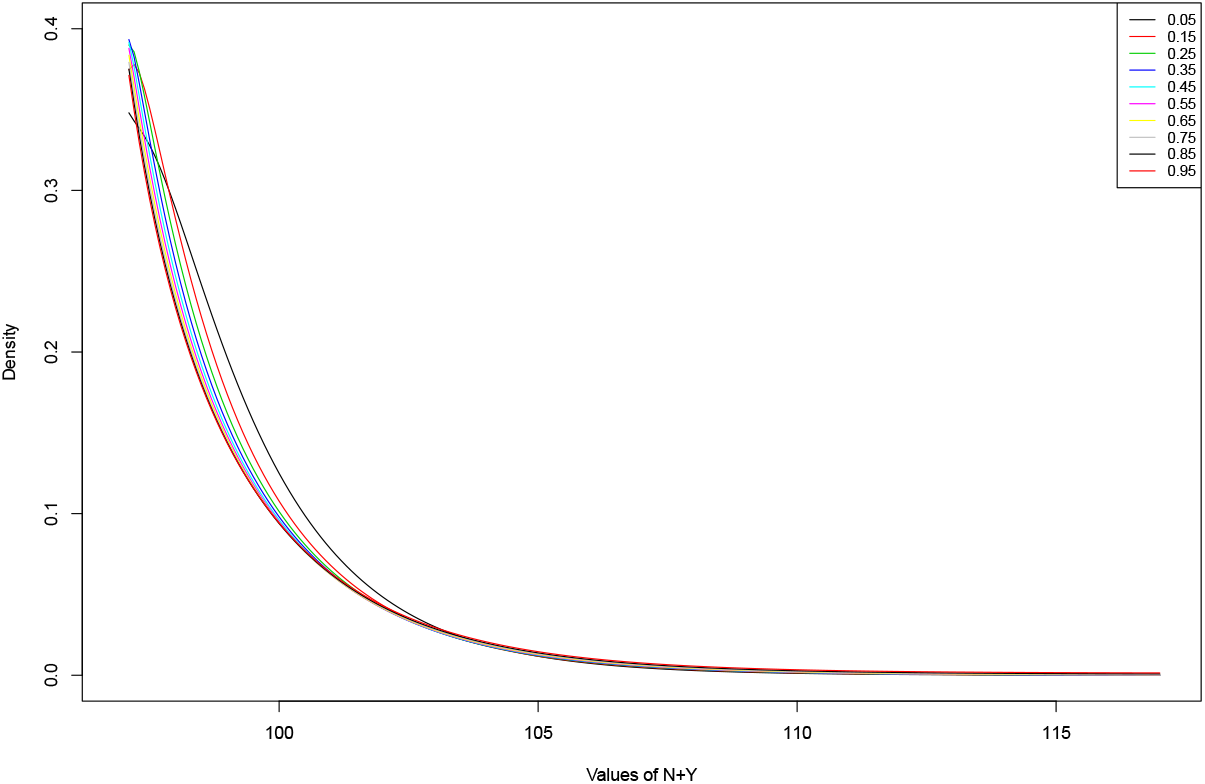
*The density* (5) *for the 1893 cohort of females*: *θ* = 2.82, *N* = 97. *Mixing function κ*(*v*) = *κ* × *v^β^ with κ* = 0.158, *β* = 7; *various α* in (0,1).

The conditional expectation can be calculated from (4) as *E*(*Y*|*V* = *v*) = *θ*(*κ*(*v*) + 1)^-1^, showing that, as *κ*(*v*) increases (*β* > 0), the expected lifetime decreases. Thus persons with higher values of V have shorter lifetimes, on average.

The cdf corresponding to the density (5) is given by integrating *V* out of (3), and from it the excess lifetime *y_p_* such that at most 100*p*% persons live beyond *N* +*y_p_*, can be obtained as a quantile. Such a value depends on 4 things: the functional form *κ*(*v*) specifying how *κ* depends on *v*; the distribution assumed for *v*; and the transition and scale parameters *N* and *θ* occurring in the STLT function. Uncertainties in specifying or estimating these quantities will be transmitted to uncertainty in derived quantities such as *y_p_*.

The operation of the model can be envisaged as follows. Drawing the lifetime of an individual, *i*, at random from the population corresponds to deciding on a value of *N*, then selecting a realized value *v* of *V_i_* from a Gamma(1/*α*, 1/*α*) distribution. This person’s lifetime is then finite, equal to *N* + *Y*, where *Y* has distribution given by (4), with a corresponding *κ*(*v*) (equivalently, a corresponding *γ*) as initially specified.

## Is there an upper limit to human lifetimes?

Much commentary on this issue related to Dong et al.^5^, some critical, some supportive, followed in *Nature* and elsewhere soon after its publication. Authors Rootzén and Zholud^12^ were critical; they rejected the inferred finite limit to human life, and, using longevity data from Japan and Western countries, suggested instead an exponential model for lifetimes beyond 110 years. Their negative criticism revolved mainly around the statistical techniques employed^5^. In response Milholland, Dong and Vijg^13^ pointed out a number of shortcomings in the analyses and conclusions of Rootzáen and Zholud^12^.

As in Huang et al.^9^, Rootzen and Zholud fitted the GPD, i.e., Eq. (1), but in their case, to data from 4 areas (North and South Europe, North America and Japan), and to a very limited set (a total of 566 life lengths) from extreme ages only (supercentenarians, 110+). In all but one instance (Japan) the estimated *γ* in their GPD fits is negative, but in no case is it significantly different from 0 (Table 5)^12^. From this Rootzén and Zholud concluded that *γ* should be taken as zero, indicating an exponential distribution for the data, and go on to argue from this for an unlimited human lifespan.

But their analysis obtaining negative *γ*, but not significantly so, is really quite consistent with that of Huang et al.^9^, where the estimated *γ* are strongly negative. The difference can be explained by the fact that only limited data from extreme ages is used in Rootzén and Zholud’s analyses, whereas in^9^ a much more extensive data set spanning ages from 65 on was used, thus obtaining greater statistical power in a demographically-informed model. A follow-up study found, almost exclusively, negative estimates of *γ* in an extensive set of Australian data (to be reported separately).

Other followup commentary occurred in Brown^14^, Hughes & Hekimi^15^, Rozing, Kirkwood & Westendorp^16^ and Lenart & Vaupel^17^. Most of those writers argued for the contrary conclusion – for no upper limit to human lifespan. Other similar criticisms are in Newman and Easteal^18^.

By contrast, a number of papers which also include discussion or estimation of *γ* have come to similar conclusions as in our paper^5,9^. We list some of them as: Alves, Neves & Rosário^19^ (they also observed no trend in supercentenarian longevity); Gbari, Poulain, Dal & Denuit^20^ (a study of 46,66 Belgians > 95 years old, finding a limit to lifetimes of ≈ 120 – 130 years and no trend across cohorts); Genz, Feifel & Pauly^21^ (a study combining the IDL and HMD databases, finding a limit of ≈ 125 – 130 years); Davison^22^; Ferriera & Huang^23^ augmented the IDL dataset with the GRG dataset and found a finite endpoint; Segers^24^; Zhou^25^; Einmahl, Einmahl & de Haa^26^ (a study of 285,000 Dutch people finding a limit of ≈ 115 – 130 years, with no trend over time) These papers generally find negative *γ*, or support its implications.

In another study, de Beer, Bardoutsos & Janssen^27^, using a logistic model, argued that an individual may live to 125 years within this century. Their model does not explicitly test the existence of a finite endpoint but tacitly assumes none exists. In other words, *γ* is not a parameter of their model; the assumptions of their model are inconsistent with a *γ* < 0, but the findings of their model cannot be used to argue in favour for or against a negative *γ*.

We can summarise the extreme value approach and its findings as follows:

- Extreme value theory suggests the GPD as a plausible distribution for extreme ages;
- At least two studies obtained good fits of the GPD to extensive data^26,9^;
- Almost all estimates of *γ* published so far, including in^12^, are negative.
- Almost all evaluations of a trend in *γ*, including^12^, found no trend over time. A detailed investigation of cohort trends in a GPD model found no trend in multiple parameters, including *γ* and *ω* for either gender^9^.

## The late-life mortality plateau and deceleration effect

Barbi, Lagona, Marsili, Vaupel & Wachter^28^ tracked every person born in Italy between 1896 and 1910 who lived to age 105 or beyond; a total of 3,836 individuals (3,373 women and 463 men). Their hazard rate analysis suggested a “mortality hazard rate plateau” between ages 105 and 110 and a consequent “late-life mortality deceleration effect” – that is, that the rate of increase in mortality with advanced age becomes slower than exponential (as it is in the Gompertz law) between those ages. This effect is usually understood to imply a model with a potentially infinite life span. The existence of a late-life mortality hazard rate plateau is also argued by Greenwood and Irwin^29^ and Gavrilov and Gavrilova^30^.

For other commentary, see Beltran-Séanchez, Austad and Finch^31^, Newman^32^ and Olshansky and Carne^33,34^. We discuss the plateau and deceleration effects in the context of our analyses below.

## Advanced age mortality accelerations and decelerations

In the STLT model fitted to the Netherlands data, a late-life mortality deceleration is indeed observed, but, remarkably, in conjunction with the previously mentioned finite upper limit to the lifespan. These conclusions appear to be inconsistent but the apparent paradox vanishes when we notice that the STLT model also implies an *advanced age mortality acceleration* rate following the late life mortality deceleration^9^. This results in estimated finite limits to the life spans.

A formula for the age *x_c_* before which the force of mortality (the hazard) under the GPD increases slower than that of the Gompertz is derived in Huang et al.^9^. After age *x_c_*, the hazard for the GPD increases faster than that of the Gompertz. In the Netherlands data, the deceleration is observed subsequent to the threshold age *N* for all cohorts and for both males and females. The ages *x_c_* at which this occurs are listed in Table 2.

**Table 2:** Estimated advanced age mortality acceleration age *x_c_*. Females: 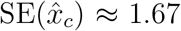 years. Males: 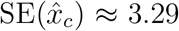 years.

| Cohort | Females | Males | Cohort | Females | Males |
| --- | --- | --- | --- | --- | --- |
| 1893 | 104.37 | 98.48 | 1901 | 102.13 | 99.24 |
| 1894 | 103.82 | 99.23 | 1902 | 105.21 | 106.79 |
| 1895 | 100.40 | 97.64 | 1903 | 107.20 | 105.12 |
| 1896 | 102.76 | 98.71 | 1904 | 104.24 | 97.93 |
| 1897 | 100.76 | 98.48 | 1905 | 106.81 | 101.97 |
| 1898 | 104.79 | 100.61 | 1906 | 106.92 | 105.53 |
| 1899 | 103.18 | 99.03 | 1907 | 104.31 | 101.16 |
| 1900 | 100.50 | 103.47 | 1908 | 102.16 | 103.31 |

## The three laws of biodemography

The *compensation law of mortality*^35,36,37^ states that for a given species, differences in death rates between different sub-populations (cohorts, in our case) decrease with age – because higher or lower initial death rates are accompanied by a lower or higher rate of mortality increase with age, such that mortality rates for different cohorts tend to equalise after some high age.

This tendency exists in the Netherlands data and is captured in the STLT model fit^9^. The threshold ages *N* at which the convergence takes place are around 98 years for females and (with somewhat greater variability) around 96 years for males (Table 1). These are close to the estimate of around 95 years made by Gavrilov and Gavrilova^36^ on quite different grounds.

The three *laws of biodemography* have been stated as:

- the Gompertz-Makeham law of mortality;
- the compensation law of mortality, and
- the existence of a late-life mortality deceleration.

The Netherlands data and the STLT model fitted to it are consistent with the compensation law of mortality, so all three laws are satisfied. The model also allows for, and for this data, predicts, a finite limit to human life span. To our knowledge, the model described here is unique in possessing all of these properties. The conclusions of the model therefore represent the most biologically and statistically rigorous evaluation of late-life human mortality to date.

## Author Contributions

All authors contributed collaboratively to the development of the ideas underlying the mixture model, and approved the final version for submission.

## Notes

### Competing Interest Statement

The authors have declared no competing interest.

